# Craniofacial shape transition across the house mouse hybrid zone: implications for the genetic architecture and evolution of between-species differences

**DOI:** 10.1101/039743

**Authors:** Luisa F. Pallares, Leslie M. Turner, Diethard Tautz

## Abstract

Craniofacial shape differences between taxa have often being linked to environmental adaptation, e.g. to new food sources, or have been studied in the context of domestication. Evidence for the genetic basis of such phenotypic differences to date suggests that within- as well as between-species variation has an oligogenic basis, i.e. few loci of large effect explain most of the variation. In mice, it has been shown that within-population craniofacial variation has a highly polygenic basis, but there are no data regarding the genetic basis of between-species differences. Here, we address this question using a phenotype-focused approach. Using 3D geometric morphometrics, we phenotyped a panel of mice derived from a natural hybrid zone between *M. m. domesticus* and *M. m. musculus*, and quantify the transition of craniofacial shape along the hybridization gradient. We find a continuous shape transition along the hybridization gradient, and unaltered developmental stability associated with hybridization. This suggests that the morphospace between the two subspecies is continuous despite reproductive isolation and strong barriers to gene flow. We show that quantitative changes in genome composition generate quantitative changes in craniofacial shape; this supports a highly polygenic basis for between-species craniofacial differences in the house mouse. We discuss our findings in the context of oligogenic versus polygenic models of the genetic architecture of morphological traits.

## Introduction

Data regarding the genetic basis of between-species differences has accumulated rapidly over the last two decades. In 1992, there were ten studies addressing “truly adaptive” traits between species pairs (Orr and Coyne 1992). By 2001, 22 new studies had addressed the genetic basis of “ordinary” between-species differences (Orr 2001). Orr defined such differences as those not involved in blocking gene flow, in contrast with sterility/viability-related phenotypes (Orr 2001). The latest and most extensive review of the “loci of evolution” includes 114 studies reporting between-species differences (Martin and Orgogozo 2013).

Sufficient data has now accumulated to enable informed discussion regarding the genetic nature of between-species differences. Long-standing questions include the role of small- vs. large-effect loci, regulatory vs. coding changes, single locus vs. many loci, and so forth. The data have shown that all such scenarios have occurred, and there is not a single or general way in which between-species differences have evolved (Orr 2001; Stern and Orgogozo 2008; Martin and Orgogozo 2013). However, some patterns seem to emerge, indicating that the evolutionary process is to some extent predictable (Stern 2013).

Speciation geneticists have been particularly interested in finding the genetic basis of postzygotic barriers resulting in sterility or reduced viability in hybrids (Orr 2001; Wolf *et al.* 2010). Given the direct relevance such traits have in reproductive isolation between species, it is understandable why “ordinary” traits have often been of secondary interest. A good example of such “ordinary” differences is craniofacial morphology. This trait has been extensively studied from a developmental perspective, for example, there are 922 protein coding genes with reported craniofacial phenotypes in the mouse (Mouse Genome Informatics-MGI database, queried on the 14.01.2016). However, the genes underlying natural variation in craniofacial traits between and within species are almost completely unknown. With the exception of *Bmp3* in dogs (Schoenebeck *et al.* 2012), and *Ptch1* in cichlids (Roberts *et al.* 2011), other data regarding between- or within-species craniofacial differences are limited to mapping studies where the causal genes underlying quantitative trait loci (QTL) still need to be identified.

The available mapping studies provide information about the genetic architecture of phenotypic differences. Results from QTL studies in fish and dog breeds suggest that few loci of large effect explain most of the craniofacial differences between species/breeds (Albertson *et al.* 2003b; Kimmel *et al.* 2005; Boyko *et al.* 2010; Roberts *et al.* 2011; Schoenebeck *et al.* 2012). In contrast, studies of within-species variation in mice have shown the opposite picture, craniofacial variation is associated with many loci of small effect (Pallares *et al.* 2015). This discrepancy has been reported for other traits where the genetic factors underlying within- and between-species variation are not the same (Orr 2001; Stern and Orgogozo 2008).

The study of craniofacial variation in mice has been a very active field in the last decades. Several studies have explored craniofacial shape variation between subspecies of the house mouse (*Mus musculus*) around the world (Macholán 2006; Boell and Tautz 2011; Siahsarvie *et al.* 2012). The effect of hybridization on craniofacial traits has been studied using wild hybrid mice (Alibert *et al.* 1994; Auffray *et al.* 1996a; Mikula and Macholán 2008; Mikula *et al.* 2010), as well as wild-derived inbred lines representing different subspecies (Renaud *et al.* 2009, 2012). Our previous study mapping craniofacial traits in hybrid zone mice identified loci which potentially contribute to within and/or between subspecies variation (Pallares *et al.* 2014). However, to date, there are no studies explicitly addressing the genetic basis of craniofacial differences between the subspecies of the house mouse. In contrast, the genetic factors underlying differences between traditional inbred lines (i.e. LG/J, SM/J, and B6) have been extensively studied (Cheverud *et al.* 1997; Leamy *et al.* 1999; Klingenberg *et al.* 2004; Wolf *et al.* 2005; Burgio *et al.* 2009; Boell *et al.* 2011; Maga *et al.* 2015).

Here, we quantify the transition of craniofacial shape along a hybridization gradient between *Mus musculus musculus* and *Mus musculus domesticus*. For this, we used first-generation lab-bred offspring of mice caught in the Bavarian region of the hybrid zone. The house mouse European hybrid zone runs from Denmark to Bulgaria marking a climatic divide between Atlantic and Continental climate. *M. m. musculus* and *M. m. domesticus* have been in contact in the hybrid zone for ∼3000 years (reviewed in Baird and Macholán (2012)).

We test the prediction that craniofacial shape differences between the two subspecies have a highly polygenic basis, *i.e.*, shape differences are caused by many loci of small effect. If this is true, we expect quantitative differences in the mean phenotype corresponding to quantitative changes in mean genome composition, and therefore: 1) the phenotypic transition should be continuous, 2) the mean shape at arbitrary points of the hybridization gradient should be different, and 3) shape changes between arbitrary points of the gradient should be directional.

## Methods

### Mouse samples

Wild hybrid mice were collected in the Bavarian region of the hybrid zone in Germany, and brought to the Max Planck Institute for Evolutionary Biology in Plön, Germany (Turner *et al.* 2012). The mice were bred in the laboratory, and first generation offspring were raised under standardized conditions. First-generation mice were sacrificed by CO_2_ asphyxiation between 9 and 13 weeks (see Turner *et al.* (2012) for details on animal experiments and ethics). 249 mice were used in this study; a subset of them were previously used in genome-wide association studies of craniofacial (Pallares *et al.* 2014) and sterility traits (Turner and Harr 2014). The sample used in this study includes siblings and half-siblings; details on the mice used in this study, including family information are presented in supplementary Table S1.

### Genotypes

Mice were classified in nine hybrid groups based on genomic background, defined as the percentage of *M. m. domesticus* alleles (Table 1). Group 0 contains individuals with 0 to 9% *M. m. domesticus* alleles, and group 9 contains individuals with 90 to 100% *M. m. domesticus* alleles. The sample lacks mice with 70 to 79% of *M. m. domesticus* alleles, and therefore hybrid group 7 was not analysed in this study. The percentage of *M. m. domesticus* alleles was estimated by the average percentage of *M. m. domesticus* alleles of the parents. In a previous study, the parents were genotyped for 37 SNPs diagnostic of the two subspecies of house mouse, *M. m. musculus* and *M. m. domesticus* (Turner *et al.* 2012). The group assignment based on average-parent value was validated for a subset of mice using estimates of percentage *M. m. domesticus* alleles based on genotypes of 270 diagnostic autosomal SNPs (Turner *et al.* 2011). Group classifications based on estimates for individuals were highly correlated with classifications based on average-parent values (r = 0.94, N= 178).

**Table 1.**
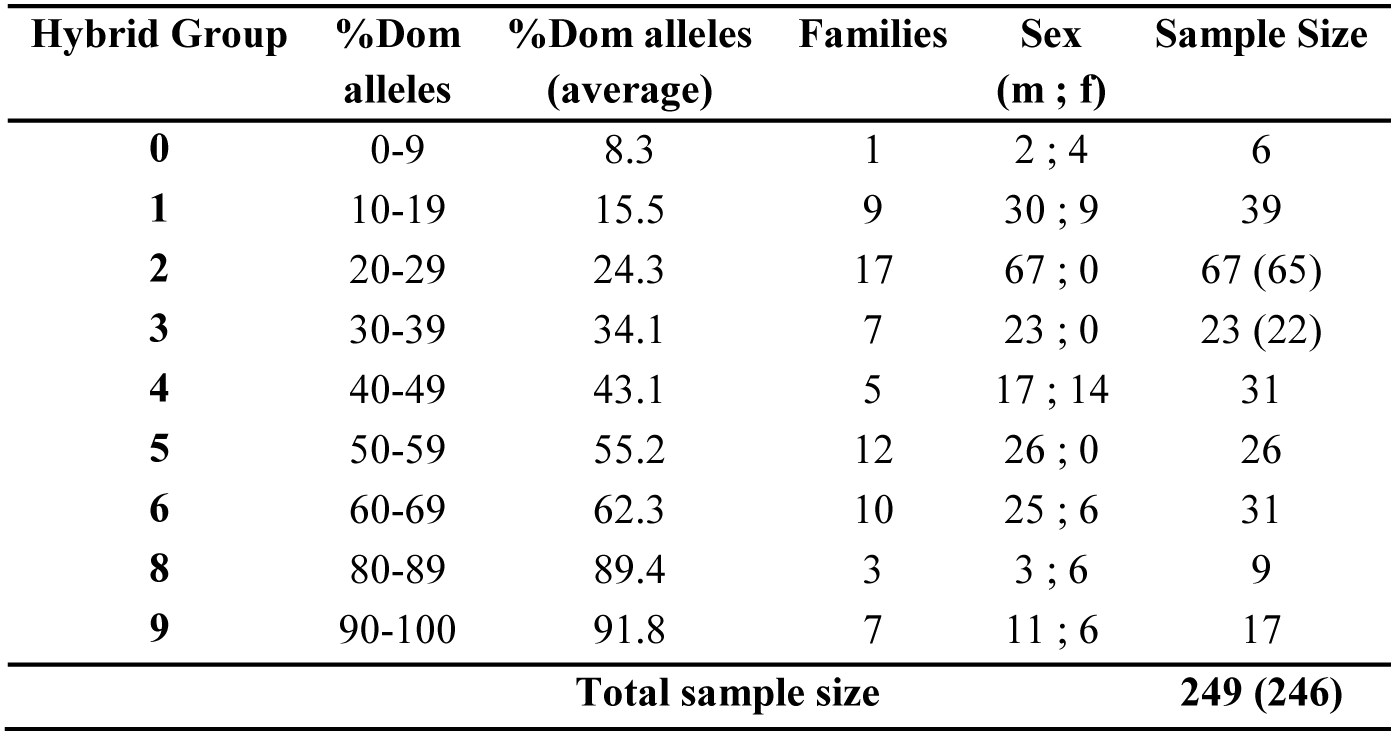
Hybrid individuals used in the analyses. Groups are defined based on the percentage of *M. m. domesticus* alleles (%Dom alleles). The number of individuals, sex, number of families, and average percentage of *M. m. domesticus* alleles per group are shown. Where sample size differs between skull and mandible, skull sample size is indicated in parentheses.

| Hybrid Group | %Dom alleles | %Dom alleles (average) | Families | Sex (m ; f) | Sample Size |
| --- | --- | --- | --- | --- | --- |
| 0 | 0-9 | 8.3 | 1 | 2 ; 4 | 6 |
| 1 | 10-19 | 15.5 | 9 | 30 ; 9 | 39 |
| 2 | 20-29 | 24.3 | 17 | 67 ; 0 | 67 (65) |
| 3 | 30-39 | 34.1 | 7 | 23 ; 0 | 23 (22) |
| 4 | 40-49 | 43.1 | 5 | 17 ; 14 | 31 |
| 5 | 50-59 | 55.2 | 12 | 26 ; 0 | 26 |
| 6 | 60-69 | 62.3 | 10 | 25 ; 6 | 31 |
| 8 | 80-89 | 89.4 | 3 | 3 ; 6 | 9 |
| 9 | 90-100 | 91.8 | 7 | 11 ; 6 | 17 |
| Total sample size |  |  |  |  | 249 (246) |

### Phenotypes

Heads were scanned using a micro-computer tomograph - microCT (vivaCT 40, Scanco, Bruettisellen, Switzerland). Energy and current were set to 70kVp and 177 μA; medium resolution was used generating a voxel size of 21 μm. 3D cross-sections were reconstructed and imported into TINA tool (Schunke *et al.* 2012) where 44 3D-landmarks were located in the skull, and 13 3D-landmarks were located in each hemimandible.

A subset of 178 mice were previously phenotyped for craniofacial shape (Pallares *et al.* 2014). However, to ensure that all mice were phenotyped under the same conditions, all 249 mice used in this study were phenotyped in the following way: Landmarks were placed using the semi-automatic landmarking tool implemented in TINA tool (Bromiley *et al.* 2014) using a reference database of ten manually-landmarked mice. Landmark position was revised and manually adjusted when necessary. Description of the landmarks used in this study can be found in Suppl. Table 2.

**Table 2.**
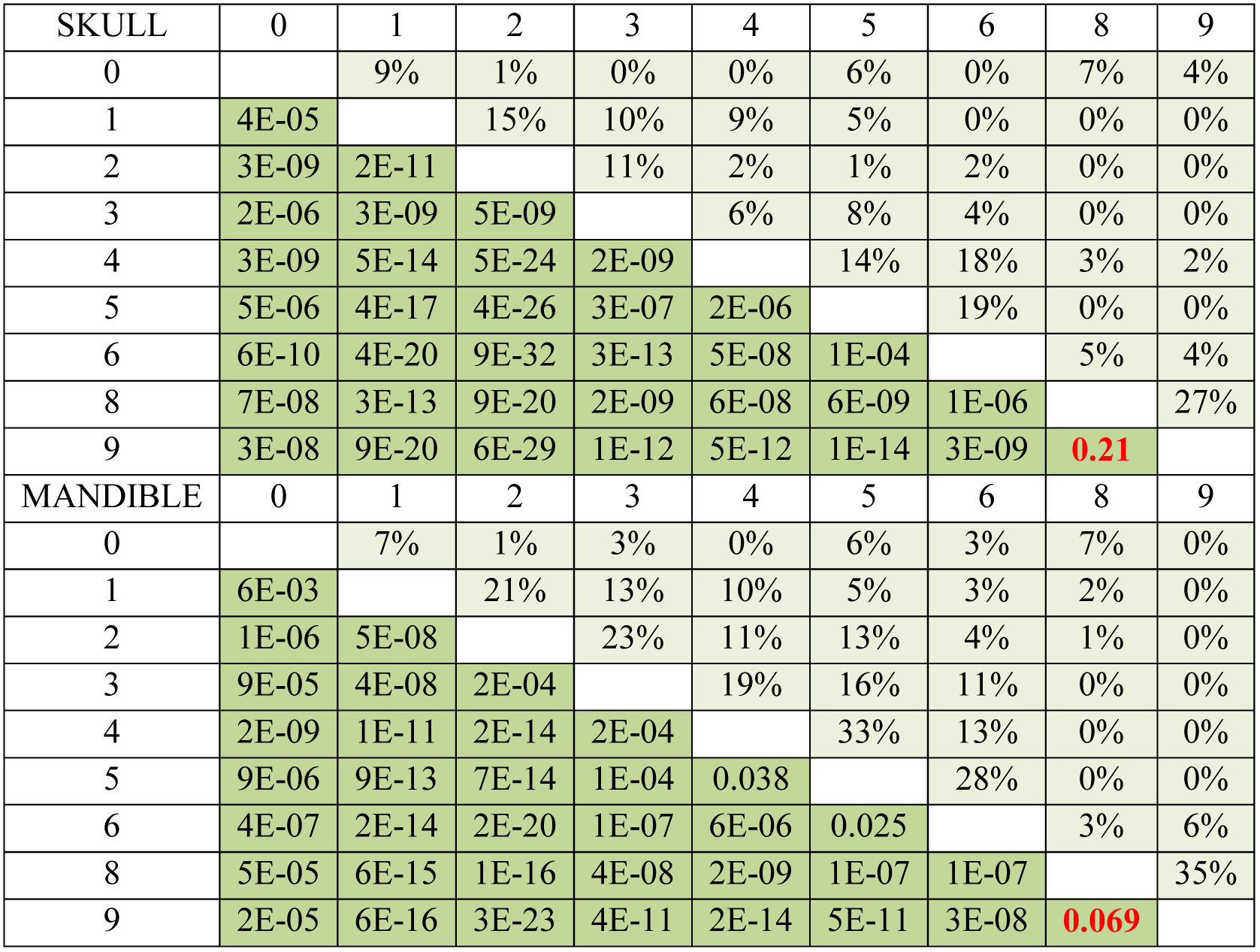
Mean shape comparisons between hybrid groups. Skull and mandible results are shown. The lower triangle (dark green) shows the p-value resulting from MANOVA. The upper triangle (light green) shows the percentage of misclassified individuals derived from a linear discriminant analysis using a leave-one-out cross-validation procedure. In red are the pairwise comparisons for groups with similar mean shape.

| SKULL | 0 | 1 | 2 | 3 | 4 | 5 | 6 | 8 | 9 |
| --- | --- | --- | --- | --- | --- | --- | --- | --- | --- |
| 0 |  | 9% | 1% | 0% | 0% | 6% | 0% | 7% | 4% |
| 1 | 4E-05 |  | 15% | 10% | 9% | 5% | 0% | 0% | 0% |
| 2 | 3E-09 | 2E-11 |  | 11% | 2% | 1% | 2% | 0% | 0% |
| 3 | 2E-06 | 3E-09 | 5E-09 |  | 6% | 8% | 4% | 0% | 0% |
| 4 | 3E-09 | 5E-14 | 5E-24 | 2E-09 |  | 14% | 18% | 3% | 2% |
| 5 | 5E-06 | 4E-17 | 4E-26 | 3E-07 | 2E-06 |  | 19% | 0% | 0% |
| 6 | 6E-10 | 4E-20 | 9E-32 | 3E-13 | 5E-08 | 1E-04 |  | 5% | 4% |
| 8 | 7E-08 | 3E-13 | 9E-20 | 2E-09 | 6E-08 | 6E-09 | 1E-06 |  | 27% |
| 9 | 3E-08 | 9E-20 | 6E-29 | 1E-12 | 5E-12 | 1E-14 | 3E-09 | 0.21 |  |
| MANDIBLE | 0 | 1 | 2 | 3 | 4 | 5 | 6 | 8 | 9 |
| 0 |  | 7% | 1% | 3% | 0% | 6% | 3% | 7% | 0% |
| 1 | 6E-03 |  | 21% | 13% | 10% | 5% | 3% | 2% | 0% |
| 2 | 1E-06 | 5E-08 |  | 23% | 11% | 13% | 4% | 1% | 0% |
| 3 | 9E-05 | 4E-08 | 2E-04 |  | 19% | 16% | 11% | 0% | 0% |
| 4 | 2E-09 | 1E-11 | 2E-14 | 2E-04 |  | 33% | 13% | 0% | 0% |
| 5 | 9E-06 | 9E-13 | 7E-14 | 1E-04 | 0.038 |  | 28% | 0% | 0% |
| 6 | 4E-07 | 2E-14 | 2E-20 | 1E-07 | 6E-06 | 0.025 |  | 3% | 6% |
| 8 | 5E-05 | 6E-15 | 1E-16 | 4E-08 | 2E-09 | 1E-07 | 1E-07 |  | 35% |
| 9 | 2E-05 | 6E-16 | 3E-23 | 4E-11 | 2E-14 | 5E-11 | 3E-08 | 0.069 |  |

Raw landmark coordinates were exported to MorphoJ (Klingenberg 2011). Raw coordinates of right and left hemimandible were subjected to a generalized Procrustes analysis (GPA) where variation due to size, position, and location is removed keeping only variation due to shape and allometry. The resulting Procrustes coordinates were averaged between right and left hemimandibles. Because the skull has object symmetry, raw coordinates were reflected and a GPA was done with the original and reflected datasets. Procrustes coordinates of original and reflected datasets were averaged per mouse. The average of right and left hemimandibles, and original and reflected skulls represent a dataset that contains information only about the symmetric component of shape. The dimensionality of skull vectors is estimated by dim = 3*k* +2*l* − 4, where *k* is the number of paired landmarks, and *l* is the number of midline landmarks (*k*=17, *l*= 10, dim = 67) (Klingenberg *et al.* 2002). The dimensionality of mandible vectors is dim = 3k – 7, where k is the number of landmarks (k = 13, dim = 32) (Klingenberg *et al.* 2002).

The asymmetric component of shape was not the main interest of this study; however, to explore developmental stability along the hybridization gradient, a value of fluctuating asymmetry (FA) was generated for each individual in MorphoJ, following Klingenberg and Monteiro (2005). Each FA value represents the individual deviation from the mean FA of the group, in units of Procrustes distance.

Centroid size (CS) was calculated for each mouse skull using all landmarks (paired and midline). Since allometry accounted for 5.4% of total shape variation, and 23% of principal component 1 (PC1), its effect was excluded from the data using the residuals of a multivariate regression of shape on CS for further analyses. CS for mandible was calculated as the average of right and left hemimandibles. Allometry accounted for 1.8% of mandible variation, and 6% of PC3. Its effect was excluded from the data following the same procedure used for skull. Differences in age have a negligible effect on shape variation (r^2^(mandible) = 0.6%, p-value=0.08; r^2^(skull) = 0.7%, p-value= 0.02).

### Size variation

Estimates of centroid size are not strongly affected by small sample size (Cardini and Elton 2007). Therefore, the mean CS of mandible and skull were calculated in MorphoJ for group 0 and 9, and these values were used as proxy for *M. m. musculus* and *M. m. domesticus* size, respectively.

To explore the variation in size across the hybrid gradient between *M. m. musculus* and *M. m. domesticus*, CS values were exported from MorphoJ into R (R-Core-Team 2015), and a linear regression of CS on hybrid group was performed. Pairwise Mann-Whitney tests were done for all pairwise comparisons of hybrid groups, and the results were adjusted for multiple testing using Holm-Bonferroni correction (R function *pairwise.wilcox.test()*).

### Shape differences between M. m. musculus and M. m. domesticus

Group 0 and 9 represent the extremes of the hybridization gradient, with an average of 8.3% and 91.8% *M. m. domesticus* alleles, respectively (Table1). However, since the estimation of mean shape is very sensitive to small sample size (Cardini and Elton 2007), groupes 0 and 1 (mus), and groups 8 and 9 (dom) were pooled to get more robust estimates of mean craniofacial shape in the extremes of the hybridization gradient.

The group *mus* contains 45 individuals with an average of 11.9% *M. m. domesticus* alleles (min 8.3%, max 18.3%). The group *dom* contains 26 individuals with 90.6% *M. m. domesticus* alleles on average (min 88.3%, max 99.3%). The mean shapes of *mus* and *dom* were generated in MorphoJ, and were used as proxy for *M. m. musculus* and *M. m. domesticus* mean shapes, respectively.

The sample size of both groups together (71 individuals) is larger than mandible shape dimensionality (dim = 32) and slightly larger than skull shape dimensionality (dim = 67). However, to be consistent with the analyses performed with smaller sample sizes (see next section), we followed Evin *et al.* (2013), and Jombart *et al.* (2010), and reduced the dimensionality of shape data by performing a principal component analysis (PCA) using all 71 mice (R function *prcomp()*) and retaining the first 10 PCs representing 79% of the total phenotypic variation in mandible shape, and 87% in skull shape.

The first 10 PCs for each trait were used in a multivariate analysis of variance (MANOVA) to assess the differences in mean shape between *mus* and *dom* (R function *aov()*). The same PCs were used in a discriminant analysis (DA) to evaluate the separation between groups. The reliability of DA was assessed by a leave-one-out cross-validation procedure (R function *lda(CV=T)*).

### Shape variation along the hybrid gradient

A multivariate regression of shape on percentage of *M. m. domesticus* alleles was used to evaluate whether skull and mandible shape was correlated with genomic admixture. The significance of the correlation was tested by permutation as implemented in MorphoJ; individuals assigned to each hybrid group were randomized without replacement and the sum of squares predicted by the regression was recorded; this procedure was repeated 10,000 times to create a null distribution. A univariate score was calculated for each mouse by projecting the shape vector of each individual onto the regression vector, in this way a visual representation of the multivariate regression was generated.

As a second approach to explore shape transition along the hybrid gradient, a between-group PCA was performed using the group means (R function *bga(type = “pca”)*).

Phenotypic variance per hybrid group was obtained in MorphoJ. It is calculated as the average of the sum of squared landmark deviations from the mean shape after Procrustes superimposition.

### Shape differences between hybrid groups

With the aim of determining if changes in average genome composition result in distinct mean craniofacial shapes, skull and mandible mean shapes were compared between groups. This resulted in 36 comparisons. Sample size for four of these pairwise comparisons is smaller than mandible dimensionality (dim = 32), and the sample size of 26 comparisons is smaller than skull shape dimensionality (dim = 67). Therefore, we reduced the dimensionality of shape data as described above to be able to compare mean shapes. In short, following Evin *et al.* (2013), and Jombart *et al.* (2010), a PCA was performed for each pair of groups, and the first 10 PCs were retained and used in MANOVA and DA. DA was validated by the leave-one-out procedure. The total variance represented by the first 10 PCs ranged from 73% to 97% depending on the pair of groups (suppl. Table 3 and 4). The significance levels resulting from MANOVA were adjusted for multiple testing using the Holm-Bonferroni method post hoc (R function *p.adjust(method=“holm”)*). The sample size of the comparison between groups 0 and 8 was 15, and therefore it was not possible to use 10 PCs in the MANOVA since it exceeded the degrees of freedom. Therefore, only in this case, 5 PCs were used in the MANOVA and DA.

Procrustes distance between groups was calculated in MorphoJ and represents shape distance. It is well known that Procrustes distances are overestimated when sample size is small, therefore, group comparisons involving group 0 (N=6), and 8 (N=9) should be interpreted with caution. Genomic distance between groups is estimated as the difference between mean percentage of *M. m. domesticus* alleles.

### Comparison of vector directions

The angle between vectors can be used to assess the similarity between vector directions (Klingenberg and McIntyre 1998; Klingenberg and Marugan-Lobon 2013). To determine if the shape transition between groups is directional, i.e. whether it resembles the overall transition between *M. m. musculus* and *M. m. domesticus*, vectors between pairs of hybrid groups were calculated and compared to the vector going from *mus* (groups 0 and 1) mean shape to *dom* (groups 8 and 9) mean shape.

First, the vector of shape transition between hybrid groups was calculated such that the groups were always compared in ascending order in terms of the percentage of *M. m. domesticus* alleles. That is, the shape vector was constructed from the group with lower percentage to the group with higher percentage (e.g. group 0 → group 1, group 2 → group 5).

Second, the vectors of shape transition between groups were compared to the vector representing shape changes from *mus* to *dom*. For this analysis, the angle between vectors was calculated and used to estimate the probability that the two vectors had the same direction due to chance. The assessment of significance was done using the formula from Li (2011) implemented in MorphoJ; and the resulting p-values were adjusted for multiple testing using the Holm-Bonferroni method post hoc.

## Results

We assigned 249 mice to 9 hybrid groups according to the percentage of *M. m. domesticus* alleles (Table1). These groups represent a hybridization continuum, starting with an average of 8.3% *M. m. domesticus* alleles (group 0) and ending with an average of 91.8% (group 9). The difference between consecutive hybrid groups is, on average, 10.4% *M. m. dom*esticus alleles, with a minimum of 2.4% between groups 8 and 9, and a maximum of 27.1% between groups 6 and 8.

There was significant sexual dimorphism in skull shape and size, however the effect on total variation is small (r^2^(shape) = 1.94%, p(10,000 permutations) = <0.0001; r^2^(CS) = 3.9%, p(10,000 perm) = 0.0019). A similar pattern was found for mandible shape and size (r^2^(shape)=2.3%, p(10,000)<0.0001); r^2^(CS)=2.15%, p(10,000)=0.0208). Despite these small differences, we pooled males and females within each group for further analysis, because the main goal of this study is to explore shape and size transitions due to genome composition, and females are relatively well distributed among the hybrid groups (see Table 1).

### Craniofacial size

There is a weak but positive correlation between hybrid group and skull CS (r^2^=2%, p-value=0.014, Figure1a), and between hybrid group and mandible CS (r^2^=1.2%, p-value=0.042, Figure1b). However, none of the pairwise comparisons of CS between groups were significant after multiple testing correction, including the extremes of the hybridization gradient (groups 0 and 9).

**Figure 1.**
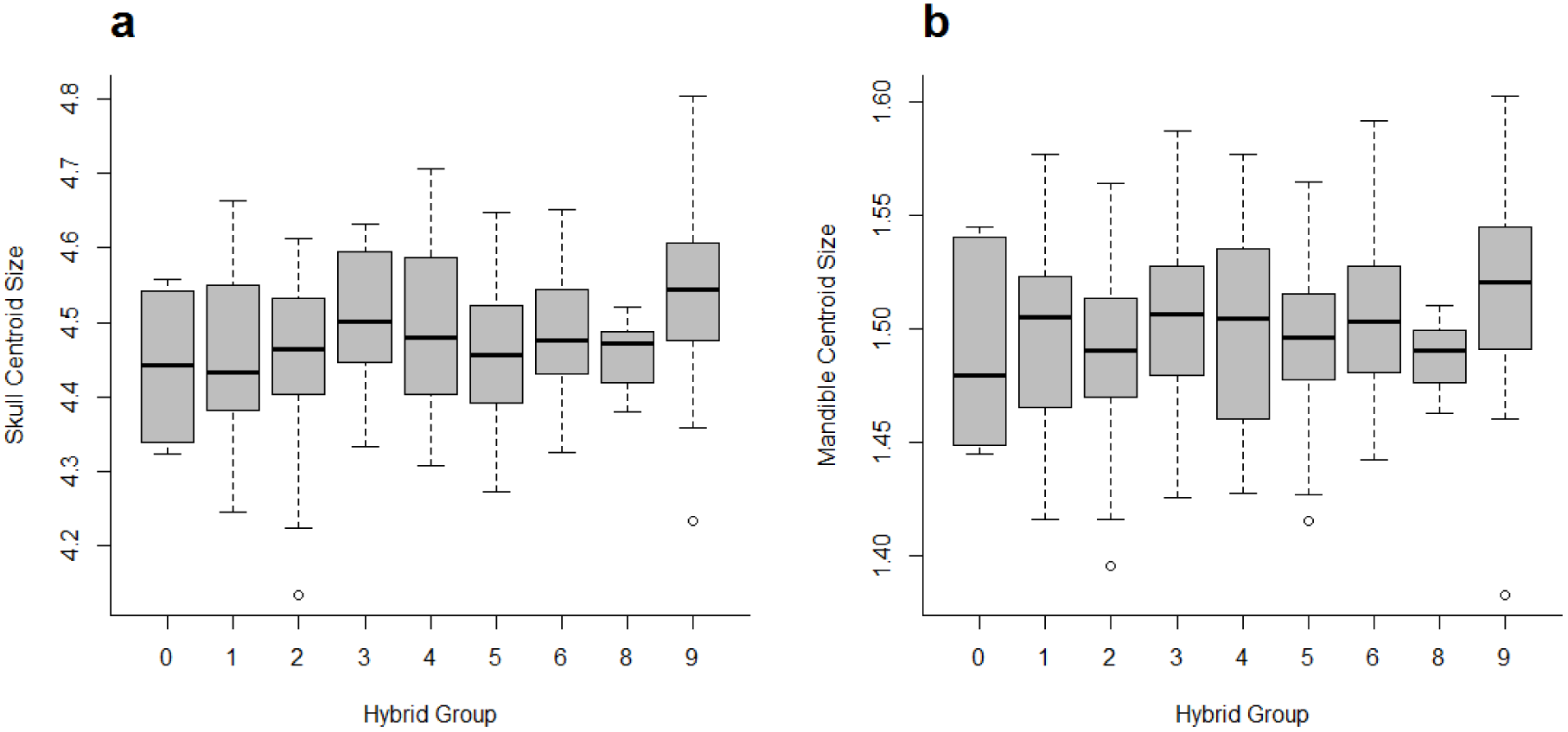
Box plot of centroid size (CS) per hybrid group. Plot (a) shows data for skull CS, and (b) for mandible CS. The box limits indicate the first and third quartile; the thick black line indicates the median CS. Linear regression of skull CS on hybrid group: p-value=0.0076, r^2^=2.5%. Linear regression of mandible CS on hybrid group: p-value=0.0438, r^2^=1.2%.

### Shape differences in the extreme of the hybridization gradient

Individuals from groups 0 and 1 (*mus*), and from groups 8 and 9 (*dom*) were combined (see Methods) and used to estimate skull and mandible shape in the extremes of the hybridization gradient. The shape differences between *mus* and *dom* were significant for skull (p-value(MANOVA) = 5×10^−25^) as well as for mandible (p-value(MANOVA) = 2×10^−26^). Individuals representing *M. m. domesticus* are characterized by straighter mandible compared to the mean *mus* mandible; this is visible by the relative arrangement of the condyle, coronoid and angular processes (Figure 2b). The lower molar row of *mus* is shorter and more distant from the body of the mandible. *Mus* individuals have also a shorter alveolar ramus, but a higher ascending ramus and a more pronounced angular process relative to the condyle (Figure 2a-c). *M. m. musculus* is characterized by a wider and higher rostrum relative to the back of the skull; shorter frontal bone relative to the nasal and parietal bones; and an upper molar row shifted towards the interior of the mouth, while individuals representing *M. m. domesticus* have a straight molar row (Figure 2d-g).

**Figure 2.**
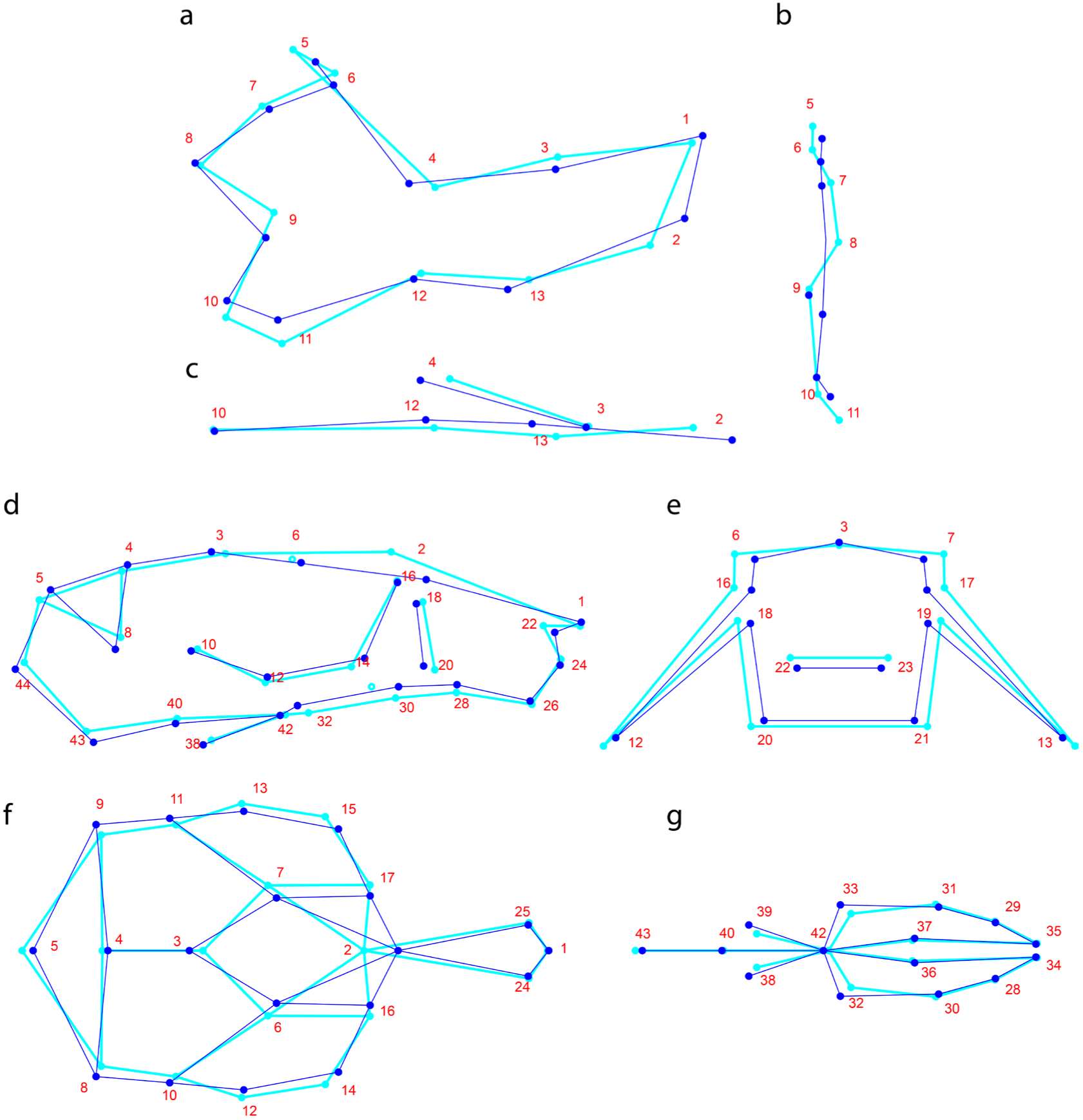
Skull and mandible shape of *mus* and *dom* groups. Light blue corresponds to the mean shape of *mus* (groups 0 and 1 combined), and dark blue to the mean shape of *dom* (groups 8 and 9 combined). Landmarks are numbered and represented as dots. 2D wireframes are used to represent the 3D shape of mandible and skull lateral view (a, d), frontal view (b, e), dorsal view (c, f), and skull ventral view (g). Shape changes are magnified 3x.

The Procrustes distance between mean skull shapes is 0.026, and 0.042 between mean mandible shapes (p(1,000 perm)<0.0001). The leave-one-out cross-validation procedure correctly assigned all individuals to their original group.

### Shape transition along the hybridization gradient

Skull and mandible shape is significantly correlated with the relative genome composition of the individuals. A multivariate regression of shape on % *M. m. domesticus* alleles showed that 8.4% of skull shape variation (p(10,000 perm) = 0.0001), and 10.8% of mandible shape variation (p(10,000 perm)=<0.0001) can be explained by average genome composition (Fig. 3). Moreover, the shape transition along the hybridization gradient is continuous, with overlaps between individual shapes found in each group, resulting in no major gaps between groups.

**Figure 3.**
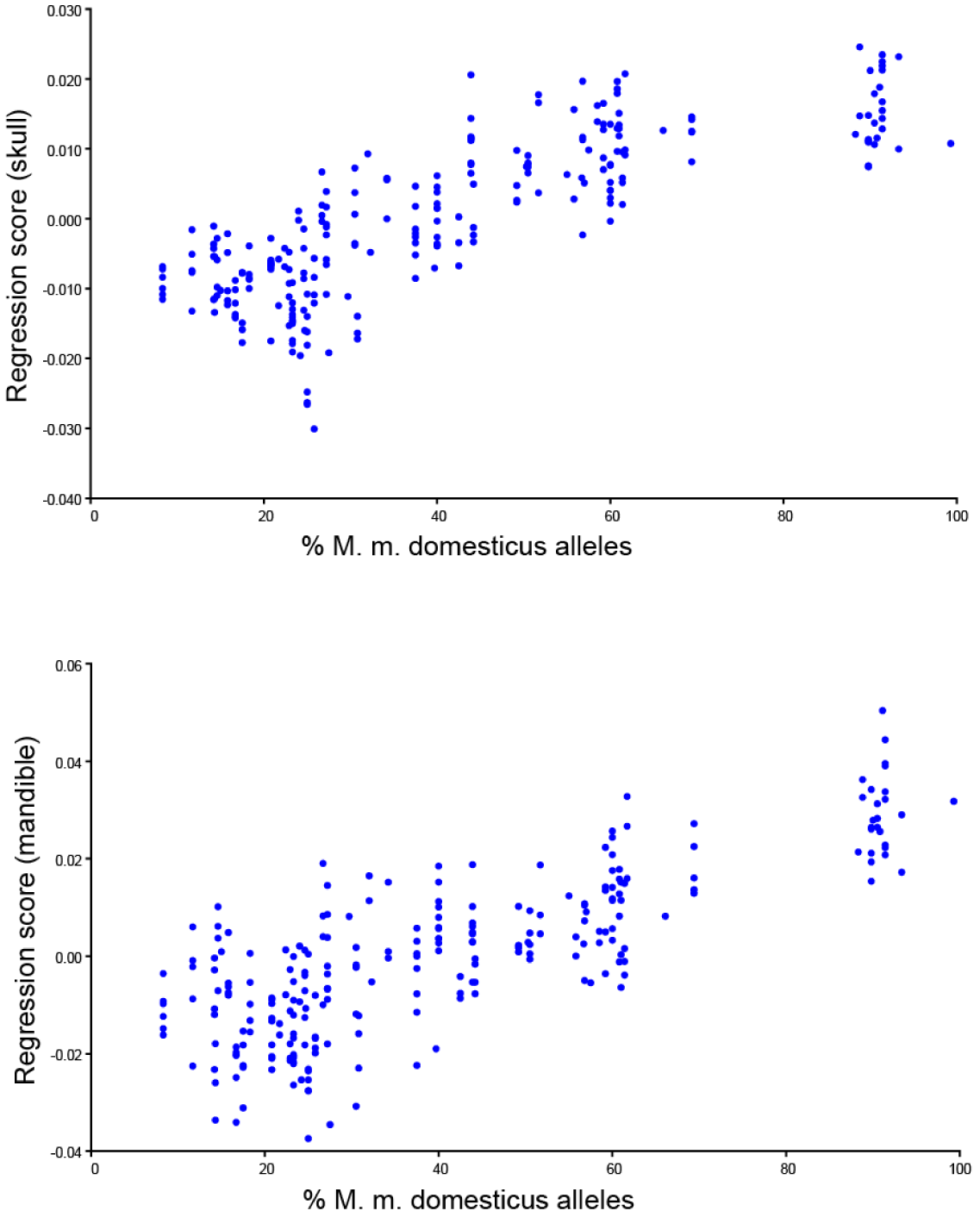
Multivariate regression of shape on genome composition. Plot (a) shows data for skull, and (b) for mandible. For the purpose of visualization a univariate score was generated for each mice. Each dot represents the projection of an individual shape vector onto the vector derived from the multivariate regression (see Methods). Skull p-value(10,000 permutations)<0.0001, r^2^=8.4%. Mandible p-value(10,000 permutation)<0.0001, r^2^=10.8%.

Continuous variation is also reflected in the strong correlation between Procrustes distance and genomic distance between groups (p-value(skull)=1×10^−8^, r^2^=61%; p-value(mandible)=2×10^−15^, r^2^=84%) - the larger the genomic distance, the larger the phenotypic distance (Figure 5a,b). However, for pairs of groups with genomic distance less than 20%, the correlation is no longer significant (Figure 5c,d).

Phenotypic variance differs between groups, but it is not correlated with the degree of hybridization (p-value(skull) = 0.72; p-value(mand) = 0.93) (Figure 4). However, variance is correlated with the number of families per group (r^2^ = 0.41, p-value = 0.036); groups with more unrelated individuals tend to have higher phenotypic variance. Skull variance ranges from 0.020 to 0.031 units of Procrustes distance, while mandible variance ranges from 0.026 to 0.040. Despite these marked changes in phenotypic variance, levels of fluctuating asymmetry remain constant across groups (Figure 4); FA in skull and mandible is, on average, 0.01 and 0.028 units of Procrustes distance, respectively.

**Figure 4.**
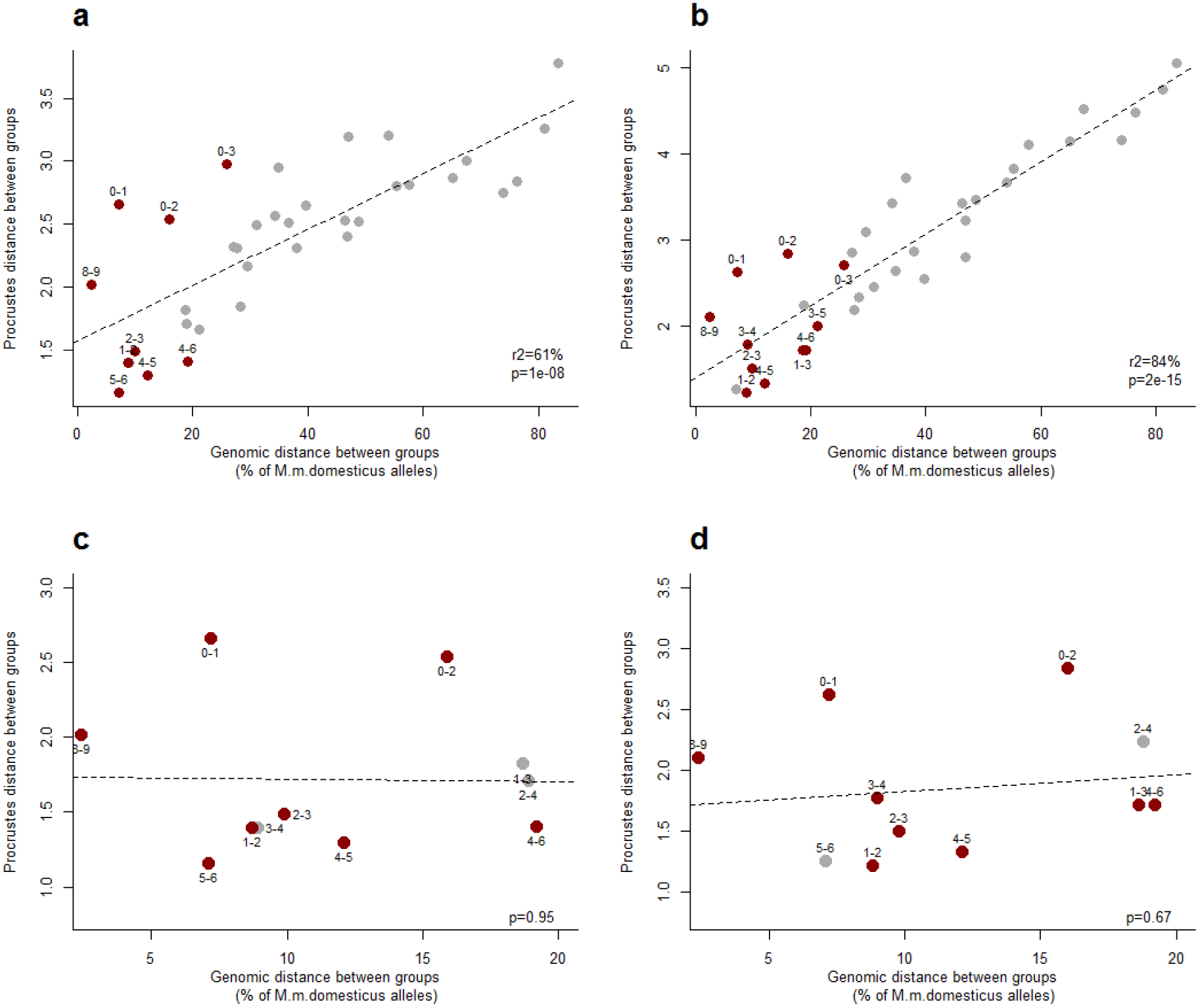
Correlation between shape distance and genomic distance. Procrustes distance between group mean shapes is used as a measure of shape distance. The difference between mean percentage of *M. m. domesticus* alleles is used as proxy for genomic distance. Each dot represents a pairwise comparison between groups. All comparisons are shown for skull (a) and for mandible (b). Pairs of groups with a genomic distance smaller than 20% are shown for skull (c), and for mandible (d). In red are pairwise comparisons without significant differences in shape (see Table 3). Procrustes and genomic distances between each pair of groups can be found in supplementary Tables S3 and S4.

### Shape differences between hybrid groups

Differences in skull and mandible mean shape were assessed for all pairs of hybrid groups. All group means differ significantly from each other, except for groups 8 and 9 (p-value(skull)=0.21, p-value(mandible)=0.07) (Table2), which showed small genomic and Procrustes distances (Figure 5).

**Figure 5.**
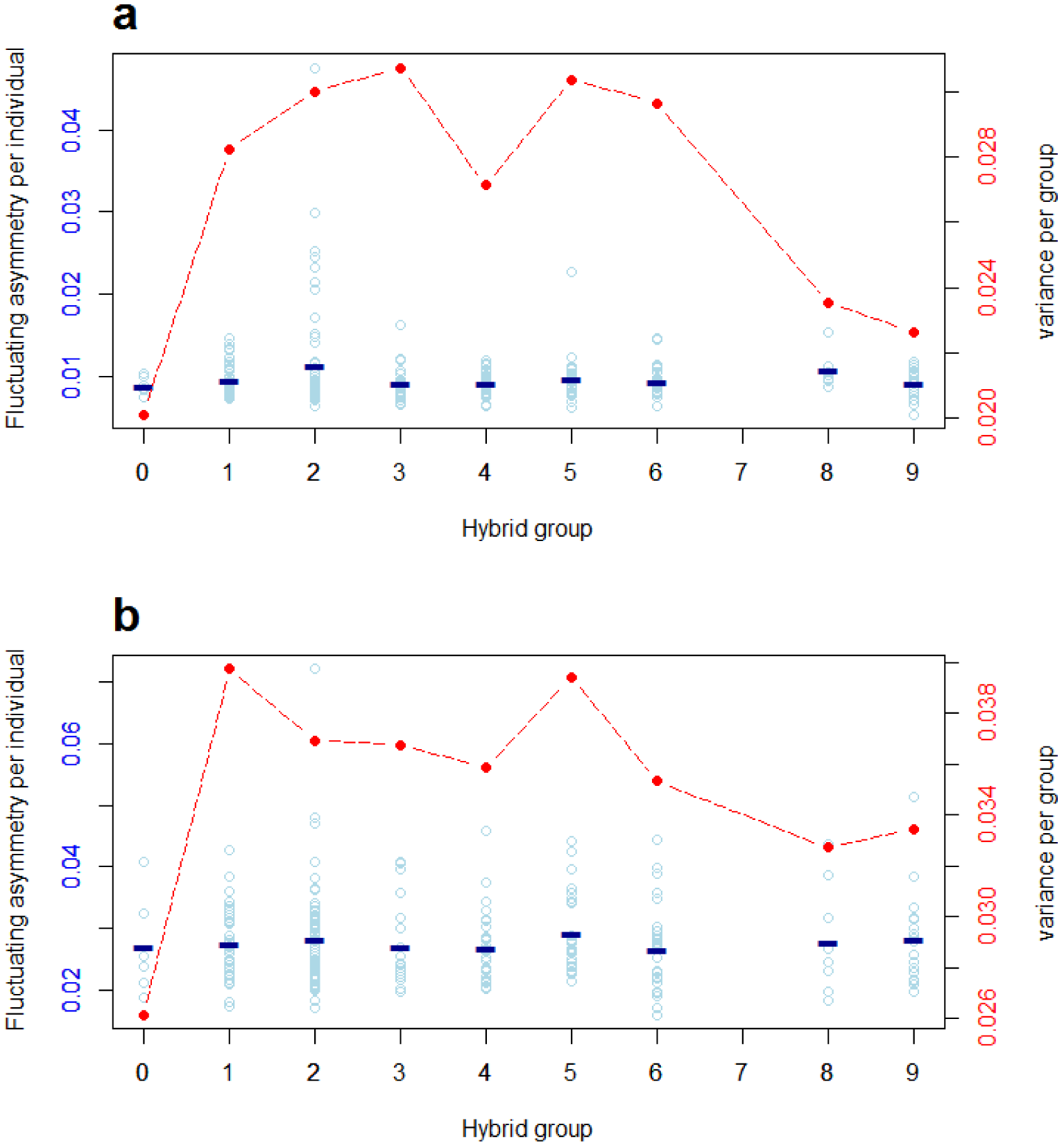
Phenotypic variation and developmental stability along the hybridization gradient. Plot (a) shows data for skull, and (b) for mandible. Fluctuating asymmetry (FA) is used as a proxy for developmental stability, FA values for individual mice are shown in blue dots; the blue line indicates the mean FA value per group. Red dots show the phenotypic variation per group, calculated as the squared root of the sum of squared landmark deviations from the group mean shape.

Figure 6 shows the ordination of the mean skull and mandible shape of all hybrid groups. A continuous transition is recovered in the first two principal component axes, with a clear gap where group 7 would likely be located if there were individuals with such genome composition in the sample of mice used here.

**Figure 6.**
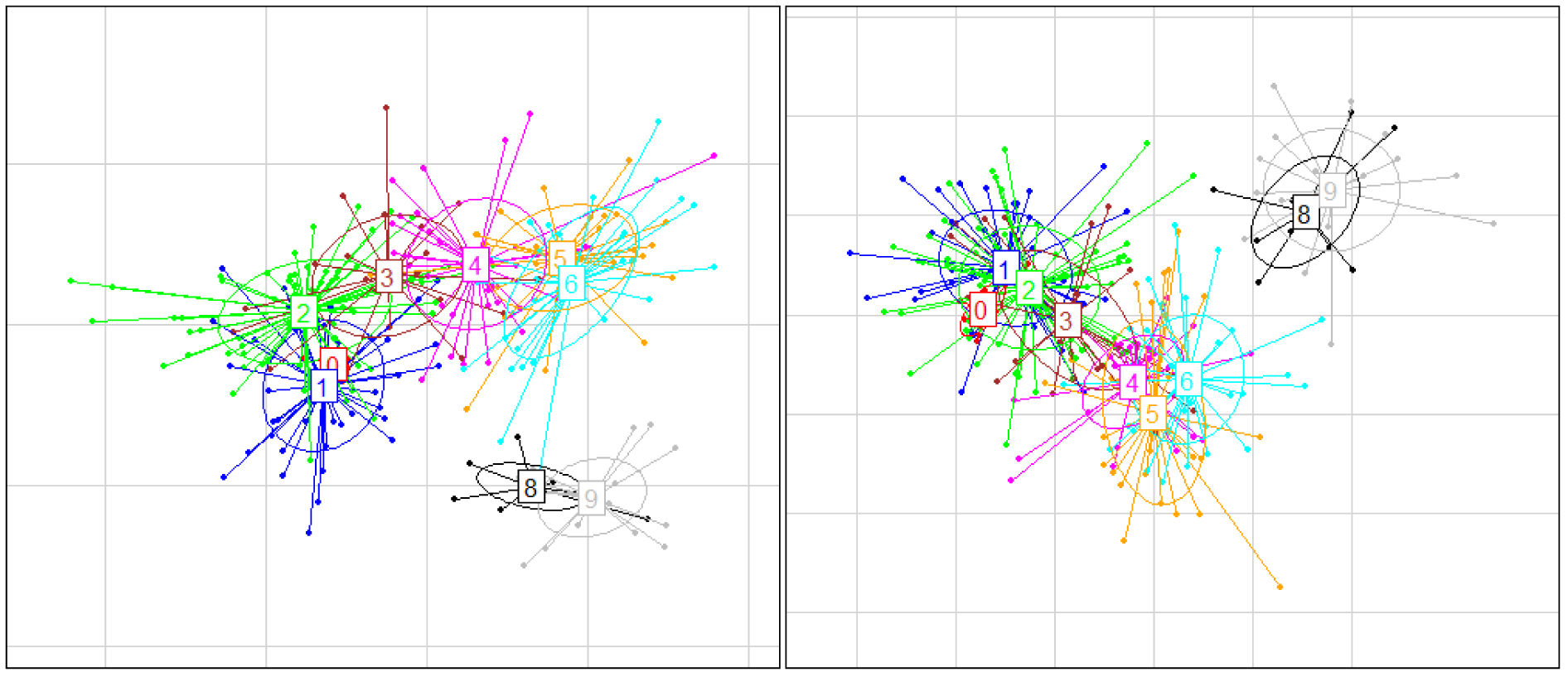
Between-group principal component analysis. The plot represents the ordination of mean skull shape (a), and mean mandible shape (b) of the hybrid groups (squares). Dots are individuals projected onto the PC axes defined by the mean shapes ordination. Each hybrid group is represented by a different colour. The numbers inside the boxes represent the groups.

### Direction of shape changes between groups

Shape change vectors were calculated between each pair of groups and compared with the shape change vector from *mus* (groups 0 and 1) to *dom* (groups 8 and 9). After multiple testing correction, 27 out of 36 pairs (75%) are consistent with the *mus*-to-*dom* skull transition, and 25 pairs (69%) are consistent with the *mus*-to-*dom* mandible transition (Table 3). Pair of groups that do not show consistent directions of change in shape space are predominantly those with small genomic distances (average distance 13.6% *M. m. domesticus* alleles), regardless of the Procrustes distance between them (Figure 5-a, b).

**Table 3.** Comparison of vector directions. The vector of shape change between groups was compared to the vector of shape change between *mus* and *dom*. Results for skull and mandible are shown. The lower triangle (dark green) shows the angle in degrees between the two vectors. The upper triangle (light green) shows the significance of the correlation between vectors after correction for multiple testing. In red are the pairs of groups whose transition along the *mus*-to-*dom* direction is not significantly supported.

| SKULL | 0 | 1 | 2 | 3 | 4 | 5 | 6 | 8 | 9 |
| --- | --- | --- | --- | --- | --- | --- | --- | --- | --- |
| 0 |  | **0.88** | **0.88** | **0.28** | 4E-03 | 4E-04 | 4E-04 | 4E-04 | 4E-04 |
| 1 | 90.3° |  | **0.83** | 0.041 | 4E-04 | 4E-04 | 4E-04 | 4E-04 | 4E-04 |
| 2 | 87.2° | 84.2° |  | **0.053** | 4E-04 | 4E-04 | 4E-04 | 4E-04 | 4E-04 |
| 3 | 78.8° | 71.0° | 72.3° |  | 9E-04 | 4E-04 | 4E-04 | 4E-04 | 4E-04 |
| 4 | 65.9° | 58.1° | 50.9° | 63.3° |  | **0.19** | **0.10** | 1E-03 | 4E-04 |
| 5 | 61.8° | 54.8° | 56.3° | 56.1° | 76.4° |  | **0.88** | 0.044 | 4E-04 |
| 6 | 60.3° | 48.2° | 50.8° | 57.0° | 74.2° | 86.2° |  | 0.044 | 4E-04 |
| 8 | 44.4° | 31.5° | 39.9° | 51.4° | 63.6° | 71.4° | 71.4° |  | **0.27** |
| 9 | 43.3° | 13.6° | 29.3° | 39.6° | 52.2° | 60.4° | 57.6° | 78.0° |  |
| MANDIBLE | 0 | 1 | 2 | 3 | 4 | 5 | 6 | 8 | 9 |
| 0 |  | **1** | **1** | **0.42** | 0.013 | 0.047 | 3E-03 | 4E-04 | 4E-04 |
| 1 | 89.1° |  | **0.99** | **0.12** | 2E-03 | 6E-03 | 4E-04 | 4E-04 | 4E-04 |
| 2 | 86.1° | 83.0° |  | **0.17** | 0.012 | 0.013 | 4E-04 | 4E-04 | 4E-04 |
| 3 | 74.8° | 67.0° | 69.8° |  | **0.13** | **0.13** | 2E-03 | 4E-04 | 4E-04 |
| 4 | 58.5° | 52.3° | 58.0° | 67.9° |  | **1** | **0.063** | 4E-04 | 4E-04 |
| 5 | 63.0° | 56.0° | 58.6° | 67.7° | 86.2° |  | 0.012 | 4E-04 | 4E-04 |
| 6 | 54.5° | 41.7° | 47.5° | 52.5° | 64.2° | 58.2° |  | 6E-04 | 4E-04 |
| 8 | 33.4° | 19.4° | 24.2° | 31.7° | 41.0° | 43.2° | 50.0° |  | **0.49** |
| 9 | 28.2° | 10.2° | 19.6° | 24.1° | 27.6° | 36.7° | 41.2° | 76.6° |  |

## Discussion

We have explored the transition of craniofacial shape and size along a hybridization gradient between two subspecies of the house mouse, *M. m. musculus* and *M. m. domesticus*. The mice used in this study are derived from wild-caught hybrids, and are thus representative of natural phenotypic and genotypic variation across the hybrid zone. At the same time, it is a sample with enhanced genetic effects relative to environmental effects because mice were raised in controlled laboratory conditions. With this design, we controlled for non-genetic factors affecting shape variation yet analyzed variation within an evolutionarily-relevant context.

### Craniofacial Size

There are no significant differences in skull or mandible size between hybrid groups. The lack of differentiation between groups 0 and 9, at the extremes of the hybridization gradient, indicates that *M. m. musculus* and *M. m. domesticus* do not differ in craniofacial size. Comparisons between wild-derived mouse strains representing the same two subspecies found the same pattern for mandible (WLA and PWK) (Renaud *et al.* 2009, 2012), and for skull (DDO and MDH) (Debat *et al.* 2000). However, differences in molar size have been reported between strains DDO and MDH (Alibert *et al.* 1997).

Heterosis of mouse mandible (Renaud *et al.* 2009), skull (Debat *et al.* 2000; Percival *et al.* 2015), and molar size (Alibert *et al.* 1997) has been reported previously, with F1 individuals being larger than the parental lines. The mice used here do not include F1 or F2 individuals, because there were no F1 hybrids sampled in the hybrid zone (Turner *et al.* 2012). Therefore, the lack of size differentiation between groups in the extreme of the hybrid gradient and the other groups indicates that any size heterosis that might have been present in the first generations of intercross between the subspecies was lost in later generations. Results from crosses between WLA and PWK inbred strains suggest that the heterosis present in F1 is lost as soon as the second generation (Renaud *et al.* 2012).

### Craniofacial Shape

In contrast with craniofacial size, skull and mandible shapes are clearly different between *M. m. musculus* (group *mus*) and *M. m. domesticus* (group *dom*). The separation between groups is perfect, even when the mice representing each subspecies have on average ∼10% alleles coming from the other subspecies. Given such clear differences between the extremes of the hybridization gradient, it is of particular interest to ask how such traits change with the percentage of *M. m. domesticus* alleles in the genome. When visually inspected, the transition along the hybridization gradient seems continuous, without major gaps between hybrid groups (Figure 3). Such a continuous pattern was first observed in a sample of wild mice collected in the Danish hybrid zone between *M. m. musculus* and *M. m. domesticus* (Auffray *et al.* 1996a). In that study, the ventral side of the skull was phenotyped using 2D geometric morphometrics. Here, we have explored the entire skull using 3D geometric morphometrics, confirming the results from Auffray *et al.* (1996b), and have expanded the analysis to the mandible, finding a similar pattern of continuous variation.

From a developmental point of view, such continual variation indicates that all craniofacial shapes derived from the random combination of the two genomes are realizable, and therefore, that the shape space between *M. m. musculus* and *M. m. domesticus* is continuous. The level of fluctuating asymmetry is constant along the hybridization gradient even when phenotypic variance changes across groups; this indicates that the degree of hybridization does not affect the developmental stability of the phenotype (Figure 4). However, results from wild-caught animals suggest that the morphological cline for ventral skull shape may be steeper than the change in hybrid index, and therefore that some impairment in skull development could be associated with certain hybrid genotypes when environmental effects and selection in the wild eare into play (Auffray *et al.* 1996a). So far, there is no formal analysis of craniofacial morphology clines in the house mouse hybrid zone. This will be necessary to estimate gene by environment effects and determine if selection is acting against some specific craniofacial morphologies.

### Shape differences between hybrid groups

A more detailed exploration of the shape transition along the hybridization gradient showed that each one of the nine hybrid groups defined based on the percentage of *M. m. domesticus* alleles has a different mean craniofacial shape, except for groups 8 and 9 that differ neither in skull nor in mandible shape. The pairwise genomic distance between groups ranges from 2.4% to 83.5% *M. m. domesticus* alleles, indicating that even small changes in genome composition generate quantifiable changes in the phenotype. It should be noted that the presence of related individuals reduces phenotypic variation within groups (see Results), however, all but one group are composed of more than one family (see Table 1), and therefore, differences in mean shape between groups cannot not be explained by family effects.

Studies in consomic lines between C57BL/6 and PWD strains (Boell *et al.* 2011), and interspecific congenic strains between C57BL/6 and SEG/Pas (Burgio *et al.* 2009; Burgio *et al.* 2012), have also shown that small differences in genome composition can generate quantifiable changes in the phenotype. However, the results from Boell *et al.* (2011) and Burgio *et al.* (2009, 2012) should be interpreted from the perspective of the individual, as inbred lines represent a single point in the universe of possible genotypes. The use of wild or outbred mice, like the sample used here, has the advantage of providing a more accurate picture of between-individual variation, because many combinations of the loci relevant for craniofacial shape are likely to be present within the same hybrid group. It therefore provides an understanding of the dynamics of the phenotype from a population perspective. Results from inbred lines showed that craniofacial shape is responsive to changes in specific loci, by contrast, our results indicate that shape is also responsive to the cumulative number of alleles at causal loci in the population.

While quantitative changes in genome composition from one group to the next generate distinct craniofacial shapes, the direction of change is conserved. Most of the pairwise comparisons between groups are consistent with differences between *M. m. musculus* and *M. m. domesticus*. This indicates that a directional walk in shape space is possible regardless of the specific loci that are being added or removed in each step, on the contrary, what seems to matter is the quantitative composition of the genome.

### Implications for the genetic basis of between-species craniofacial differences

Taken together, our results suggest that many loci of small effect determine the craniofacial differences between *M. m. musculus* and *M. m. domesticus*. The shape differences between subspecies are caused by different alleles at these loci, but the interactions among them appear predominantly additive, since they produce developmentally stable phenotypes in the hybrids; this is evident from the constant values of fluctuating asymmetry along the hybridization gradient. Further, the smooth transition between the subspecies excludes the possibility that few loci of large effect are responsible for between-species differences.

By contrast, loci of large effect have been implicated in craniofacial differences in other taxa. Between 23 and 46 QTLs have been associated with craniofacial differences between species of cichlids, with individual QTL explaining up to 52% of phenotypic variation (Albertson *et al.* 2003b, a). Later, it was shown that *Ptch1* is the causal gene underlying a large-effect locus for lower jaw differences between cichlid species (Roberts *et al.* 2011). A study of morphological variation in dogs explored 55 traits including many craniofacial measurements; the top three SNPs per trait explained on average 67% of phenotypic variation between breeds (Boyko *et al.* 2010). A different study showed that five QTL explain most of the skull variation between dog breeds, and identified *Bmp3* one of the underlying causal genes (Schoenebeck *et al.* 2012). Different stickleback ecotypes have very characteristic craniofacial shapes, one to five QTL have been found to explain these differences (Kimmel *et al.* 2005).

Although it is known that mapping studies overestimate the absolute phenotypic effect of each QTL, the overall message from these studies is that relatively few loci are responsible for within- and between-species craniofacial variation, which seems to be in contrast with our results. However, there is a major difference between these studies and ours with respect to the history of adaptation of the corresponding taxa. In the fish and dog studies, craniofacial diversity is associated with an adaptive radiation with strong selective pressures during domestication, respectively. Under such scenarios, it has been predicted that phenotypic differences are generated by an exponential distribution of mutational effects (Orr 1998), where few mutations of large effect generate an initial rapid change in the phenotype that later is refined by mutations of small effect.

Major adaptive changes in morphology are not evident in the house mouse system. Although some specific adaptations appear to occur when new habitats or islands are invaded (Renaud and Auffray 2010; Boell and Tautz 2011; Renaud *et al.* 2013; Babiker and Tautz 2015), these remain subtle. Hence, the divergence between separated mouse populations and subspecies may be equally influenced by neutral accumulation of allelic differences. Under this scenario we do not expect to see loci of large effect underlying between-subspecies differences. However, the relative importance of drift and selection still needs to be quantified.

Using a GWAS approach, we have previously shown that the genetic architecture of within-population craniofacial shape variation in the mouse is highly polygenic (Pallares *et al.* 2015). In this study, using a phenotype-focused approach, we find patterns that support a highly polygenic basis for between-subspecies differences. Different experimental approaches (GWAS, congenic lines, consomic panels) using different type of mice (inbred, outbred, wild) have found a positive correlation between the length of genomic fragments and the magnitude of their effect on craniofacial shape (Burgio *et al.* 2009; Boell *et al.* 2011; Burgio *et al.* 2012; Pallares *et al.* 2014; Pallares *et al.* 2015). Here we also find a positive correlation between genomic distance and phenotypic distance. Therefore, the evidence seems strong enough to suggest that, irrespective of the taxonomic level at which differences are being observed, craniofacial shape variation in the mouse has a highly polygenic basis.

## Conclusions

The polygenic architecture of shape determination in the house mouse results in a situation in which more or less arbitrary mixtures of different alleles result in a developmentally stable phenotype with smooth transitions between different degrees of admixture. This implies that in cases of micro-evolutionary adaptation to environmental changes, many genes could be involved in modifying the shape in the required direction.

## Acknowledgements

We thank Sabrina Renaud for fruitful discussions, and Bettina Harr for sharing mouse samples. This work was supported by institutional funds of the Max Planck Society to DT and funds from the Deutsche Forschungsgemeinschaft to B. Harr (SFB-680). This project was developed while LFP was part of the International Max Planck Research School (IMPRS) for Evolutionary Biology.

